# Functional analysis of *ESRP1/2* gene variants and *CTNND1* isoforms in orofacial cleft pathogenesis

**DOI:** 10.1101/2024.07.02.601574

**Authors:** Caroline Caetano da Silva, Claudio Macias Trevino, Jason Mitchell, Hemma Murali, Casey Tsimbal, Eileen Dalessandro, Shannon H. Carroll, Simren Kochhar, Sarah W. Curtis, Ching Hsun Eric Cheng, Feng Wang, Eric Kutschera, Russ P. Carstens, Yi Xing, Kai Wang, Elizabeth J. Leslie, Eric C. Liao

## Abstract

Orofacial cleft (OFC) is a common human congenital anomaly. Epithelial-specific RNA splicing regulators *ESRP1* and *ESRP2* regulate craniofacial morphogenesis and their disruption result in OFC in zebrafish, mouse and humans. Using *esrp1/2* mutant zebrafish and murine Py2T cell line models, we functionally tested the pathogenicity of human *ESRP1/2* gene variants. We found that many variants predicted by *in silico* methods to be pathogenic were functionally benign. *Esrp1* also regulates the alternative splicing of *Ctnnd1* and these genes are co-expressed in the embryonic and oral epithelium. In fact, over-expression of *ctnnd1* is sufficient to rescue morphogenesis of epithelial-derived structures in *esrp1/2* zebrafish mutants. Additionally, we identified 13 *CTNND1* variants from genome sequencing of OFC cohorts, confirming *CTNND1* as a key gene in human OFC. This work highlights the importance of functional assessment of human gene variants and demonstrates the critical requirement of *Esrp*-*Ctnnd1* acting in the embryonic epithelium to regulate palatogenesis.

## Introduction

The study of orofacial cleft (OFC) has been foundational to genetic analysis of congenital anomalies. Craniofacial structural malformations are amenable to detailed phenotypic classification in large cohorts where genomic studies have been carried out to identify associated loci (1–8). As whole-genome sequencing (WGS) strategies and technologies advance, a growing list of genes and gene variants associated with OFC are being cataloged (1, 8–11). These approaches have uncovered the critical role of many genes regulating the embryonic oral epithelium in palate formation and OFC pathogenesis, including: *TP63*, *IRF6*, *GRHL3*, *ESRP1/2*, *CTNND1* (12–24).

Because most cases of non-syndromic OFC occur sporadically, the pathogenicity of variants cannot be inferred or supported by segregation among affected family members. Therefore, determining the functional significance of gene variants remains challenging. Multiple *in silico* predictive algorithms such as SIFT, PolyPhen-2, MutationTaster, PROVEAN and AlphaMissense offer functional predictions for gene variants utilizing amino acid sequence information, sequence conservation, biophysical properties, or homolog alignment (25–30). However, when given the same gene variants, these predictive tools may provide null values or contradicting results (31, 32). Indeed, the American College of Medical Genetics and Genomics and the Association for Molecular Pathology (ACMG-AMP), weights functional studies higher than in silico evidence for asserting pathogenic potential in gene variants for genes not previously established as causal for a particular disease (33–35). We and others previously showed that functional testing of human gene variants is essential, as *in silico* approaches alone fail to reach the necessary accuracy for clinical translation (36–40). While bioinformatics tools have greatly facilitated the functional interpretation of genetic variants (41–43), it is also important to note the essential role of functional validation of gene variants, especially for those genes where computational predictions tend to differ from experimental validation (44–50).

*ESRP1* and its paralog *ESRP2* are epithelial splicing regulatory proteins that co-localize with *Irf6* and function in the embryonic epithelium to regulate craniofacial development and epithelial-mesenchymal transition during embryogenesis (22, 51–53). Global transcriptome analysis comparing mutant *irf6* and wildtype zebrafish revealed that the epithelial-specific splicing regulator *Esrp1* was differentially expressed (52). We showed that *Esrp1* and *Esrp2* are colocalized in the periderm and oral epithelium and are required for the formation of the anterior neurocranium (ANC), a teleost embryonic structure developmentally analogous to the mammalian primary palate in the manner that it is formed from the convergence of frontonasal derived midline prominence and paired maxillary projections (54–57). Targeted disruption of *Esrp1* in the mouse resulted in bilateral cleft lip and palate (21). In the *esrp1/2* double homozygote zebrafish, cleft formed in the ANC and extended to the upper edge of the mouth opening, analogous to the cleft lip and/or palate (CL/P) phenotype observed in the *Esrp1/2* mutant mice (22, 52). In humans biallelic *ESRP1* mutations were described to cause hearing loss (58), heterozygous *ESRP2* mutations were associated with CL/P (20) and both *ESRP1* and *ESRP2* splicing targets were related to cancer-associated processes (59). Given the central role of *ESRP1* in periderm and embryonic epithelial development, there is likely selection against deleterious *ESRP1* alleles so that variants associated with hearing deficit are likely hypomorphic and homozygous or biallelic loss-of-function alleles are likely embryonic lethal and not observed clinically.

Here, we applied complementary *in vivo* and *in vitro* models to functionally interrogate human *ESRP1* and *ESRP2* gene variants. To increase the rigor of the functional test using another independent assay, we also examined Esrp-mediated alternative splicing in a murine *Esrp1/2* double knockout Py2T cell model. The Py2T cell line has been used effectively to study epithelial mesenchymal transition and we have previously generated and characterized Esrp1 and Esrp2 double knock-out Py2T lines (23, 53). Using these independent approaches, we functionally determined the pathogenicity of the 7 *ESRP1* and 12 *ESRP2* human gene variants from CL/P cohorts or reported in hearing loss. We previously showed that *Esrp1/2* regulated splicing of *Ctnnd1* (60). Using RNAscope, we found that *Ctnnd1* transcripts co-localized with *Esrp1* and *Esrp2* in the mouse and zebrafish embryonic oral epithelium. The *esrp1/2* zebrafish model also presented a functional assay to test the function of Esrp-regulated genes such as *Ctnnd1*. In fact, exogenous expression of *ctnnd1* mRNA in zebrafish *esrp1/2* mutants partially rescued the cleft ANC, foreshortened pectoral fin and fused otolith phenotypes. Additionally, WGS of CL/P cohorts identified 13 new *CTNND1* gene variants, making this one of the most frequently associated genes in OFC. Taken together, these results demonstrate the critical requirement of *Esrp*-*Ctnnd1* operating in the embryonic epithelium to regulate palatogenesis.

## Methods

### Animal husbandry and breeding

Zebrafish (*Danio Rerio)* of the Tübingen strain were raised and bred following approved institutional protocols at Massachusetts General Hospital. Embryos were collected and raised in E3 Medium (5.0 mM NaCl, 0.17 mM KCl, 0.33 mM CaCl2, 0.33 mM MgSO4) containing 0.0001% Methylene blue at 28.5°C.

### Gene variant identification, sequence alignment, and variant effect prediction

Three WGS datasets of 759 OFC trios from the Gabriella Miller Kids First (GMKF) Research (dbGaP; European trios, dbGaP: phs001168.v2.p2; Colombian trios, dbGaP: phs001420.v1.p1; Taiwanese trios, dbGaP: phs000094.v1.p1) were filtered for variants in *ESRP1, ESRP2,* and *CTNND1* that were (1) heterozygous in the affected patient, (2) had a minor allele frequency no greater than 0.001 in any population in gnomAD or 1000 Genomes, and (3) had a variant consequence of missense, frameshift, stop-gain, splicing, or in-frame insertion/deletion. We further supplemented the resulting list with additional variants from ClinVar associated with an OFC or autosomal recessive deafness. In total, the ClinVar list included 12 *ESRP1* and 20 *ESRP2* variants. ClinVar variants were accessed in 2021, we note that new variants have been uploaded to ClinVar for ESRP1 and ESRP2, but these new variants did not include relevant clinical phenotype information so were not included in this study. OFC associated genes were based on a previously published study that curated a list of approximately 500 genes based on known clinical syndromes and association results from GWAS (61).

To further refine the variant list to identify variants for testing in mouse and zebrafish assays, we aligned the human, mouse and zebrafish *Esrp1* and *Esrp2* amino acid sequences using Clustal Omega (62). 7 *ESRP1* and 12 *ESRP2* variants at fully conserved residues were then annotated using SIFT, PolyPhen-2, and AlphaMissense to obtain the predicted change in protein function and were categorized as benign, pathogenic, or of unknown significance. We included a silent mutation from *ESRP2*, at threonine 475 (T475T) that served as an internal negative control. Variants were annotated to the following human transcripts: *ESRP1*: NM_017697.4/ ENST00000433389.8; ESRP2: NM_024939.3/ENST00000473183.7; and *CTNND1*: NM_001085458.2/ENST00000399050.10.

All variants from this study are listed in Table 1 in the supplementary material.

**Table 1:** *ESRP1* and *ESRP2* variants classification.

| ESRP1 | Protein Domain | PolyPhen-2 | SIFT | Alpha Missense | Zf in vivo assay | Py2T in vitro assay | Interpretation |
| --- | --- | --- | --- | --- | --- | --- | --- |
| Q90R |  | Damaging | Damaging, LC | Benign | Rescue | n/a | Benign |
| E194A |  | Benign | Benign | Benign | Rescue | n/a | Benign |
| D222fs |  | n/a | n/a | n/a | Mutant | Deficient | Damaging |
| L259V | RRM1 | Damaging | Benign | Pathogenic | Rescue | Restored | Benign |
| K287R | RRM1 | Damaging | Benign | Benign | Rescue | n/a | Benign |
| Y605F |  | Benign | Damaging, LC | Benign | Rescue | n/a | Benign |
| N643S |  | Benign | Benign | Benign | Rescue | Restored | Benign |

| ESRP2 | Protein Domain | PolyPhen-2 | SIFT | Alpha Missense | Zf in vivo assay | Py2T in vitro assay | Interpretation |
| --- | --- | --- | --- | --- | --- | --- | --- |
| L92Q |  | Benign | Damaging, LC | Likely pathogenic | Rescue | n/a | Benign |
| R250Q |  | Damaging | Damaging | Likely pathogenic | Rescue | Restored | Benign |
| R315H | RRM1 | Damaging | Damaging | Likely pathogenic | Mutant | Deficient | Damaging |
| R353Q | RRM1 | Damaging | Damaging | Likely pathogenic | Rescue | Restored | Benign |
| C372S | RRM2 | Benign | Benign | Ambiguous | Rescue | n/a | Benign |
| R437H | RRM2 | Damaging | Benign | Ambiguous | Rescue | n/a | Benign |
| T475T | RRM3 | Benign | Benign | n/a | Rescue | Restored | Benign |
| S508L | RRM3 | Damaging | Benign | Likely pathogenic | Rescue | Restored | Benign |
| R520STOP | RRM3 | n/a | n/a | n/a | Mutant | Deficient | Damaging |
| E547del | RRM3 | n/a | n/a | n/a | Rescue | n/a | Benign |
| L665W |  | Benign | Damaging, LC | Likely benign | Rescue | n/a | Benign |
| R667C |  | Damaging | Damaging, LC | Ambiguous | Rescue | Restored | Benign |

### Rare-variants analysis

We performed rare variant burden tests using RV-TDT (2) for protein-altering variants in *ESRP1*, *ESRP2*, and *CTNND1* that had a minor allele frequency less than 0.1% in any gnomAD population. DenovolyzeR (0.2.0), an R package which compares the observed number of DNMs to the expected number of DNMs based on a mutational model developed by Samocha et al. (2014) (63), was used to determine if de novo variants were enriched in these three genes.

### Plasmid generation, site-directed mutagenesis, and mRNA synthesis

mRNA from wildtype zebrafish embryos was collected at multiple time points from 6 hours post fertilization (hpf) to 4 days post fertilization (dpf), reverse transcribed, and combined to make pooled cDNA to clone the *esrp1* coding sequence (CDS). *esrp1* and *esrp2* were each cloned into a pCS2+8 plasmid backbone using the In-Fusion HD Cloning Kit (Clontech). The resulting pCS2+8-*esrp2* plasmid was mutagenized with synonymous mutations surrounding the translational start-site using the GeneArt site-directed mutagenesis (SDM) system (ThermoFisher) to generate *esrp2* transcripts resistant to *esrp2* morpholino binding. The 19 human *ESRP1* and *ESRP2* variants were each individually introduced to the pCS2+8-*esrp1* or MO-resistant pCS2+8-*esrp2* plasmids through the GeneArt SDM system. All generated pCS2+8 plasmids were digested with NotI at 37°C for 1hr, and capped mRNA was synthesized using the SP6 mMessage mMachine kit (ThermoFisher).

For the murine Py2T transfection experiments, we used the pIBX-C-FF(B)-mCherry-*esrp1*(2A)-+CKLP plasmid containing the mouse *Esrp1* cDNA sequence, fused to a mCherry tag (gift from Russ Carstens, University of Pennsylvania). Mouse *Esrp2* cDNA was purchased from Genomics Online. *Esrp1* cDNA was cloned into the pcDNA3.1 backbone containing a CMV promoter and SV40 polyA tailing sequence for expression in mammalian cells using the In-Fusion HD Cloning Kit (Clontech) to generate the *pcDNA3.1-esrp1-mCherry plasmid*. An mCherry tag was fused in-frame onto the *Esrp2* cDNA and introduced into the pcDNA3.1 backbone through a multi-insert in-Fusion cloning strategy, using the pIBX-C-FF(B)-mCherry-Esrp1(2A)-+CKLP as the template for the 2A-mCherry sequence to generate the pcDNA3.1-*esrp2-mCherry plasmid*. Selected human *ESRP1* and *ESRP2* gene variants were introduced using the GeneArt SDM system, as described above.

### Zebrafish microinjection and *esrp1/2* rescue assay

We previously generated a zebrafish line carrying homozygous loss-of-function alleles in *esrp1* through CRISPR/Cas9 harboring −4 bp indels which led to a frame shift mutation and early protein truncation (52). *esrp2* morpholinos (GeneTools) were reconstituted to a concentration of 8ug/uL in water and stored in single-use aliquots at RT. 2nL droplets containing (1) 8ng *esrp2* morpholino, (2) 0.05% phenol red and (3) 200pg of *esrp1, esrp2,* or *esrp* gene-variant mRNA were microinjected directly into the cytoplasm of one-cell stage *esrp1^−/−^* zebrafish embryos and grown until 4dpf. (We have previously shown that the *esrp2* morpholino, injected into *esrp1^−/−^ esrp2^wt/wt^* is sufficient to phenocopy the *esrp1^−/−^; esrp2 ^−/−^* phenotype, which is consistent with previous descriptions (22, 52). Since all the injected embryos were derived from mating of *esrp1^−/−^* males and females, all animals had the *esrp1^−/−^* genotype and did not require additional genotyping after phenotype analysis. At 4 dpf, embryos were fixed in 4% formaldehyde, stained with acid-free Alcian blue as previously described (64), and micro-dissected to inspect the anterior neurocranium (ANC). The ANC phenotype flatmount was then scored as wildtype ANC, cleft ANC or rescued ANC.

### PY2T cell maintenance and transfection

Mouse Py2T cells and *Esrp1/2* DKO Py2T cells were a gift from Russ Carstens from the University of Pennsylvania (23). Cells were maintained in DMEM supplemented with 10% FBS and penicillin/streptomycin and were not cultured past passage 30. 10.8ug of plasmid was transfected onto 10^6^ cells using the 100uL Neon system (ThermoFisher) with a single, 30 second pulse at 1400V and plated onto 6-well plates. Cells were harvested for RNA after 24hr, reverse transcribed, and the cDNA was used for RT-PCR using primers spanning the splice junctions for *Ctnnd1* exons 1 and 3 and *Afhgef11* exons 36 and 38, Arhgef11 Forward (TCAAGCTCAGAACCAGCAGGAAGT) and Arhgef11 Reverse (TGCTCGATGGTGTGGAAGATCACA), as described (23). The gels were quantified by densitometry using Fiji/ImageJ and the results are expressed as mean ± SEM. Statistical analysis involved using GraphPad Prism 9.0 for Windows. The experiments were performed in triplicate. One-way Anova test, with each comparison standing alone was used for statistical analysis. *P* < 0.05 was considered statistically significant.

### *ctnnd1* mRNA injection into *esrp1−/−; esrp2+/−* intercross

To construct the mRNA in vitro transcription (IVT) template, synthetic Ctnnd1 cDNA, isoform-201 on Ensembl (ENSDART00000106048.4), was cloned into the linearized DNA template vector (Takara Bio USA). The plasmid vectors were purified by a QIAprep spin miniprep kit (QIAGEN). The plasmid was digested with Hind III HF (NEB Biolabs) at 37°C for 1hr, 80°C for 20 minutes for inactivation and mRNA was synthesized using the T7 MEGAshortscript kit (ThermoFisher).

For micro-injection, progeny of *esrp1−/−; esrp2+/−* inter-cross, previous described by Carroll, 2020 (52) were injected at the single cell stage with either 250 pg of *ctnnd1* mRNA (along with water), or *gfp* mRNA, for controls. Injected embryos were raised to 4 dpf, at which time embryos were fixed in 4% formaldehyde, stained with acid-free Alcian blue, and microdissected to inspect the anterior neurocranium (ANC). The ANC was scored as wildtype ANC or cleft ANC. Additionally, the pectoral fins were also analyzed and scored as wildtype fin or curled fin. For the otolith phenotype, wildtype was scored when the otoliths were separate and the mutant phenotype when the otoliths were fused. For the paired bilateral structures, if one fin was curled or one set of otoliths were fused, the animal was scored as mutant. After the phenotypic assessments for ANC, fin and otoliths, both the mRNA injected embryos and the control injected embryos was tracked and individually genotyped. Whenever there is an animal with genotype of *esrp1^−/−^; esrp2^−/−^* but exhibited ANC that are not fully cleft, fins that are not fully curled and separate otoliths, these animals were scored as rescues.

### RNA in situ hybridization staining (RNAScope and BaseScope)

Wildtype and *esrp1^−/−^; esrp2^+/−^* zebrafish were crossed and the progeny embryos raised to 4 dpf. The *esrp1^−/−^; esrp2^−/−^* double mutant embryos were scored at 4 dpf based on the abrogated pectoral fin phenotype. The wild type and *esrp1^−/−^; esrp2^−/−^* embryos were fixed in 4% formaldehyde, taken through a sucrose gradient, and then cryo embedded and sectioned. RNAScope probes were designed with assistance from ACDBio to target the region of 700-1661 base pairs of the RNA for DR Ctnnd1 XM_021476936.1, which corresponds to ENSDART00000106048.4 for ensemble 201.

Additionally, RNAScope and BaseScope probes were designed for murine Esrp1 (we have previously shown that Esrp1 and Esrp2 colocalize in the oral epithelium) (52). Hybridization and staining were performed according to the manufacturers protocol. Stained sections were imaged on a Leica SP8 confocal microscope where a Z-stack was obtained and analyzed on imageJ software to obtain optimal images. BaseScope probes were designed and purchased from ACDBio to specifically target the Ctnnd1 long and short isoforms. Staining was carried out according to the manufacturer’s protocols on both fixed, frozen, and sectioned wildtype and Esrp1/2 DKO at E15. Stained sections were imaged as above.

### Statistics and Reproducibility

The results are expressed as percentage or as mean ± SEM. Statistical analysis was using GraphPad Prism 10 for Windows (GraphPad Software, San Diego, CA, www.graphpad.com). All experiments were performed at least in triplicate. Two-way analysis of variance or Student *t* test was used for statistical analysis. *P* < 0.05 was considered statistically significant.

## Results

### *esrp1* and *esrp2* are required for morphogenesis of epithelial derived tissues

We previously described the genetic requirement of *esrp1/2* in zebrafish epithelial development, disruption of which resulted in tethering of the upper mouth opening extending into a separation of the anterior neurocranium, a phenotype morphologically analogous to CL/P of amniotes (52). Given the expression of *esrp1/2* and in periderm and embryonic epithelial cells broadly, we examined other structures formed by epithelial origins. It was reported that *Esrp1* regulated the alternative splicing of *Arhgef11*, which was described to be important for proper otoliths development in zebrafish (65). When the *esrp1/2* double mutants were examined at 4 dpf, we discovered that more than 90% of the mutant larvae exhibited at least one fused otolith (Figure 1A, B).

**Figure 1.**
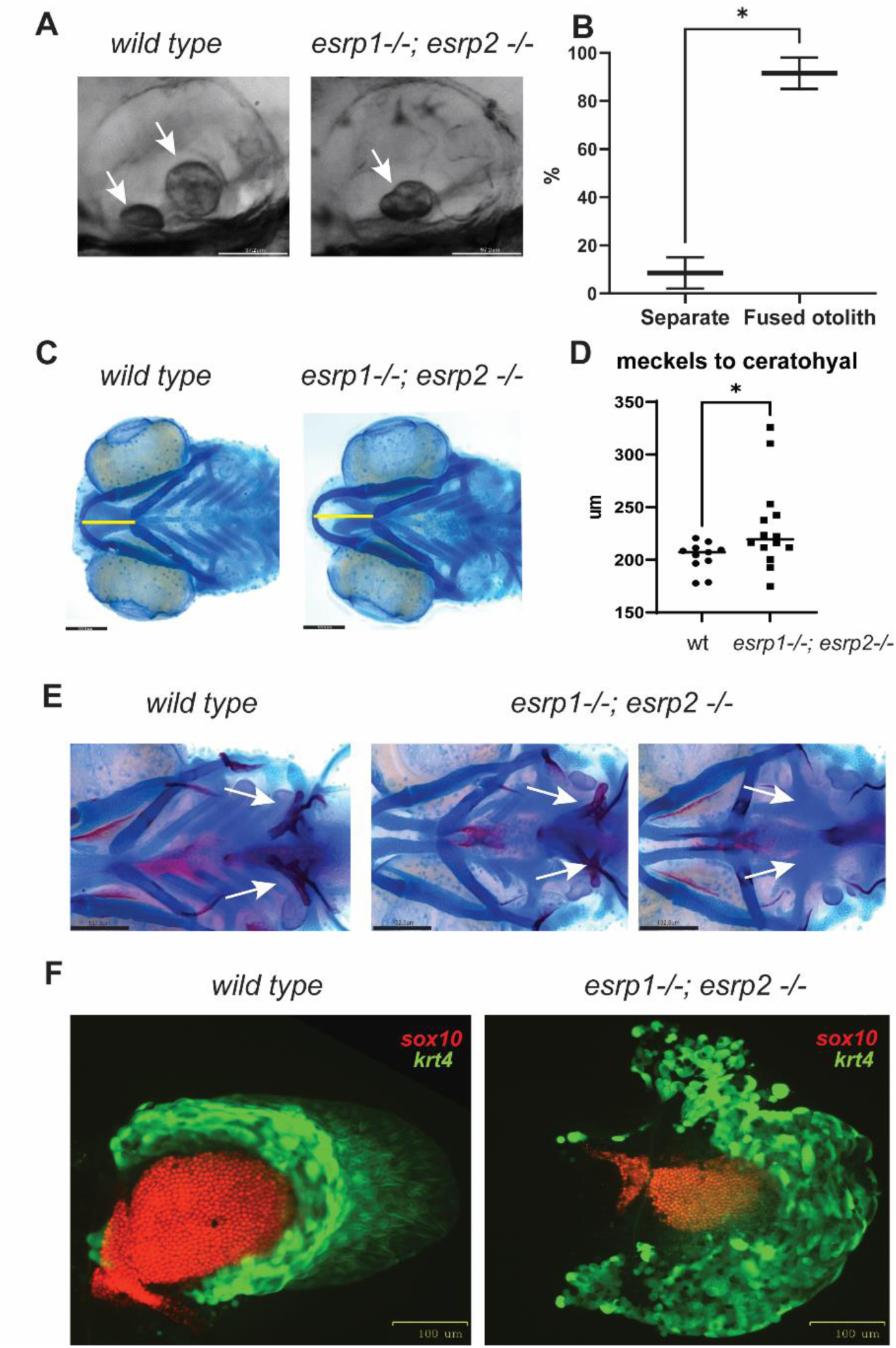
*esrp1* and *esrp2* are required for morphogenesis of epithelial derived tissue: otoliths, pharyngeal teeth and pectoral fins. **(A)** zebrafish otoliths indicated by white arrows at 72 hpf. **(B)** Quantification and t test of zebrafish otoliths from genotyped mutants characterized as separate or fused otoliths. t-test, n=75. **(C)** Alcian blue representation of a 6 dpf zebrafish wildtype and esrp1^−/−^ esrp2^−/−^ double mutant showing cartilage stain, yellow line shows the measurement of the distance between the midline of Meckel’s and ceratohyal cartilages. **(D)** quantification and t-test analysis of this measurement in wildtype (n=11) and *esrp1^−/−^; esrp2 ^−/−^* mutants (n=14). **(E)** Alcian blue and Alizarin red staining of larvae at 7 dpf ventral view, the pharyngeal teeth are present in wildtype (white arrows). In contrast, the *esrp1^−/−^; esrp2^−/−^*all exhibit decreased number of teeth, and occasionally some double mutants lack all ceratobranchial cartilages and the pharyngeal teeth are absent. **(F)** wildtype and esrp1^−/−^ esrp2^−/−^ mutant pectoral fins labeled with *sox10* mCherry (red) and *krt4* gfp (green).

Ventral cartilages that form with epithelial-mesenchymal interactions were also dysmorphic, where the Meckel’s cartilage appeared longer in the antero-posterior axis and narrower in the coronal axis. These morphologic differences can be captured by measuring the distance between Meckel’s and ceratohyal cartilages which is extended in the *esrp1/2* mutants (Figure 1C, D). We also detected partial penetrance of loss of ceratobranchial cartilages in 30% of the *esrp1/2* double mutant larvae at 7 dpf, and these larvae also exhibited loss of pharyngeal teeth (Figure 1E).

Epithelial-mesenchymal interaction is also required for pectoral fin development. We observed that the *esrp1/2* double mutants exhibit foreshortened and curled pectoral fins, where the *sox10* labeled chondrocytes that populate the mesenchymal component and the *krt4* labeled epithelial populations are both decreased in cell number in the *esrp1^−/−^; esrp2^−/−^* fins at 4 dpf (Figure 1F). Whereas the wildtype fins extend and fan out as they develop to 4 dpf, the fins in the *esrp1^−/−^; esrp2^−/−^* larvae curl proximally and are typically stuck to the torso through epithelial attachments (Figure 6A).

### *In vitro* and *in vivo* assays to functionally test *ESRP1* and *ESRP2* human gene variants

In a previous study we showed that *esrp1*−/−; *esrp2*+/− intercross yielded Mendelian ratio of 25% *esrp1*−/−; *esrp2*−/−, and that injection of morpholino against *esrp2* in the *esrp1*−/− mutant embryos can consistently phenocopy the *esrp1*−/−; *esrp2*−/− double mutant (Figure 2A) (52). This *esrp1*−/−; *esrp2* MO model provides significant advantages over *esrp1*+/−; *esrp2*+/− intercross, as the entire clutch of the *esrp1*−/− embryos injected with *esrp2* MO consistently exhibited the cleft ANC phenotype greatly facilitating detection of rescue of injected *ESRP1/2* mRNA to be tested.

**Figure 2.**
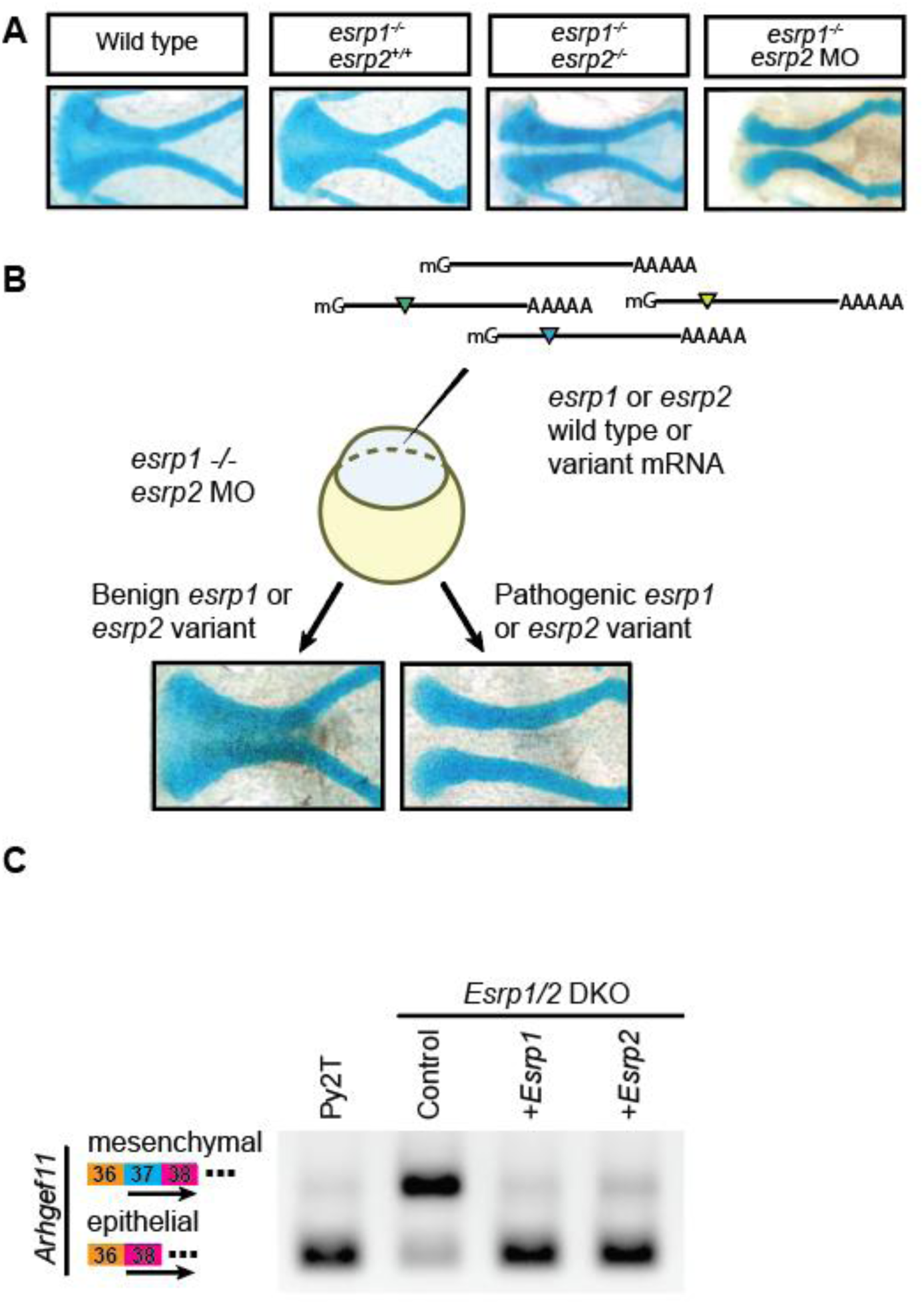
Complementary *in vivo* and *in vitro* functional assays to test human *ESRP1* and *ESRP2* gene variants. **(A)** Microdissected ANC of Alcian-blue stained embryos at 4 dpf for wild-type, *esrp1^−/−^*; *esrp2^+/+^, esrp1^−/−^; esrp2^−/−^,* and *esrp1^−/−^; esrp2 MO* embryos. **(B)** Schematic for the *esrp* morphant variant assay in zebrafish. Variants that robustly rescued the cleft ANC phenotype were scored as benign, while variants that failed to rescue the cleft ANC phenotype were scored as pathogenic. **(C)** RT-PCR was performed using primers spanning exons 36-38 of *Arhgef11* on cDNA isolated from wild-type mouse Py2T cells, *Esrp1/2* double-knockout Py2T cells, or *Esrp1/2* double-knockout PyY2T cells electroporated with plasmids encoding for either *Esrp1* or *Esrp2* genes. Arrow markers point to the epithelial (short) isoform and mesenchymal (long) isoform retaining exon 37.

We found that over-expression of wildtype zebrafish and human *ESRP1* and *ESRP2* mRNA rescued the cleft ANC phenotype in *esrp1*−/−; *esrp2* MO embryos (Figure 2B) (52). Alcian blue staining of *esrp1^−/−^*; *esrp2^−/−^* zebrafish at 4 dpf revealed a cleft ANC phenotype where a population of chondrocytes in the medial ANC is absent. A similar phenotype is observed when translation-blocking anti-*esrp2* morpholinos were injected into *esrp1^−/−^* embryos (Figure 2A).

To functionally test human *ESRP1* or *ESRP2* gene variants, we introduced point mutations into zebrafish *esrp1* or *esrp2* coding sequences and subsequently co-inject 8ng of anti-*esrp2* MO with either: (1) capped *esrp1* mRNA, (2) capped *esrp2* mRNA mutagenized with synonymous mutations at the MO binding site, or (3) either *esrp1* mRNA encoding for human *ESRP1* gene variants of unknown significance, or (MO-resistant) *esrp2* mRNA encoding for human *ESRP2* gene variants of unknown significance. We hypothesized that benign variants that preserve protein function would robustly rescue the cleft ANC phenotype like native *esrp1* or *esrp2* mRNA. Conversely, pathogenic human *ESRP1/2* gene variants with loss-of-function would fail to rescue the cleft ANC phenotype (Figure 2B). Human *ESRP1* and *ESPR2* gene variants were cloned by site directed mutagenesis, and synthesized mRNA was injected with *esrp2* MO into one-cell stage *esrp1^−/−^* embryos. The *esrp2* cDNA was engineered to prevent hybridization of the *esrp2* MO to the synthesized mRNA.

In order to gain additional functional assessment of the gene variants, we developed an independent *in vitro* assay using *Esrp1/2* mutant Py2T cells (66). The murine Py2T epithelial cell line was developed where *Esrp1* and *Esrp2* were ablated using CRISPR-mediated gene editing. The *Esrp1/2−/−* Py2T cells exhibited splicing deficiencies in the *Esrp* target gene, *Arhgef11* (Figure 2C) (66). RT-PCR performed on wildtype Py2T cell cDNA using primers spanning splice junctions for *Arhgef11* demonstrated the presence of two major isoforms. The difference between these two isoforms is the presence or absence of exon 37, which is included in mesenchymal cells, but skipped in Py2T epithelial cells (23, 67, 68). Py2T cells carrying *Esrp1* and *Esrp2* loss-of-function alleles preferentially expressed the longer mesenchymal isoform of *Arhgef11*.

We found that over-expression of *Esrp1* or *Esrp2* in the *Esrp*1/2 DKO Py2T cells efficiently rescued RNA-splicing to generate the epithelial isoform of *Arhgef11* transcript (Figure 2C).

### Identifying human *ESRP1* and *ESRP2* gene variants

Genome sequencing efforts have deposited numerous gene variants in publicly available repositories, including the Gabriella Miller Kids First (GMKF) Pediatric Research Program and ClinVar (69–71). We filtered sequencing data from the both repositories for patients with OFC or autosomal recessive deafness (20, 58) and identified gene variants for either *ESRP1* or *ESRP2* to generate a list of 32 potentially disease-associated gene variants.

Because we are utilizing *in vivo* assay in zebrafish and *in vitro* assay in murine Py2T cells, we prioritized those human *ESRP1* and *ESRP2* gene variants residing in cross-vertebrate conserved residues. For ESRP1, the overall amino acid sequence identity was 97% and 64.68% between humans and mice, or humans and zebrafish, respectively. However, when focusing on the RNA-recognition motif (RRM) domains of ESRP1, the similarity of the sequences between humans and mice and humans and zebrafish increased to 98.82% and 94.12% for RRM1, 99.08% and 79.82% for RRM2, and 95.06% and 77.78% for RRM3. Similarly, for ESRP2, the overall amino acid sequence similarity was 98.67% between humans and mice and 85.33% between humans and zebrafish. The domain-specific amino acid sequence similarities were 98.67% and 85.33% for RRM1, 98.13% and 81.31% for RRM2, and 96.3% and 77.78% for RRM3 between humans and mice, and humans and zebrafish, respectively. Altogether, we identified 19 out of the 32 gene variants in residues fully conserved between human, mouse, and zebrafish. Gene variants were evenly spread throughout both proteins and included two variants in the RRM1 domain of *ESRP1* and two variants each in the RRM1, RRM2, and RRM3 domains of *ESRP2* (Figure 1 Supplementary Material).

We found that the *in silico* predictions from SIFT and Polyphen-2 followed one of four patterns: (1) concordant predictions from both tools annotating the variant as benign, (2) concordant predictions from both tools annotating the variant as damaging, (3) discordant predictions from both tools, (4) tools unable to predict the effect of the variant on protein function (Table 1). Altogether, two variants from *ESRP1* (E194A and N643S) and two variants from *ESRP2* (C372S and T475T) were predicted by both SIFT and PolyPhen-2 to have a benign effect on protein function. One variant from *ESRP1* (Q90R) and four from *ESRP2* (R250Q, R315H, R353Q, and R667C) were predicted by both to have a deleterious effect on protein function. SIFT and PolyPhen-2 do not offer predictions for three truncation variants (*ESRP1* D222fs, *ESRP2* R520*, and *ESRP2* E547del). However, the remaining three *ESRP1* variants (L259V, K287R and Y605F) and four *ESRP2* variants (L92Q, S508L, R437H, and L665W) had discordant predictions between both algorithms. Thus, *in silico* predictions were not adequate to annotate roughly half of the selected gene variants and required an alternate approach to predict their effects on protein function.

### Functional testing of *ESRP1* and *ESRP2* variants in zebrafish and murine Py2T cell assays

The selected 19 *ESRP1* and *ESRP2* gene variants were experimentally tested in zebrafish and Py2T cell assays. Site-directed mutagenesis was carried out in *ESRP1* and *ESRP2* cDNA sequences and cloned into the pCS2+8 vector backbone to generate capped mRNA for microinjection into zebrafish embryos. The zebrafish assay was optimized by microinjection of *esrp2* translation-blocking morpholinos into *esrp1^−/−^* intercross, because the *esrp2*^−/−^ females are infertile (22). However, since the *esrp2* MO would also neutralize exogenous injected *ESRP2* mRNA upon co-injection into zebrafish embryos, synonymous mutations were introduced in the translational start site of the pCS2+8-*Esrp2* plasmid, to generate *esrp2* MO-resistant *ESRP2* mRNA transcripts. Co-injection of 8ng of *esrp2* MO with 200pg of either *ESRP1* mRNA or MO-resistant *ESRP2* mRNA fully rescued the ANC phenotype in over 75% of 19 injected clutches at 4 dpf (Figure 3A).

**Figure 3.**
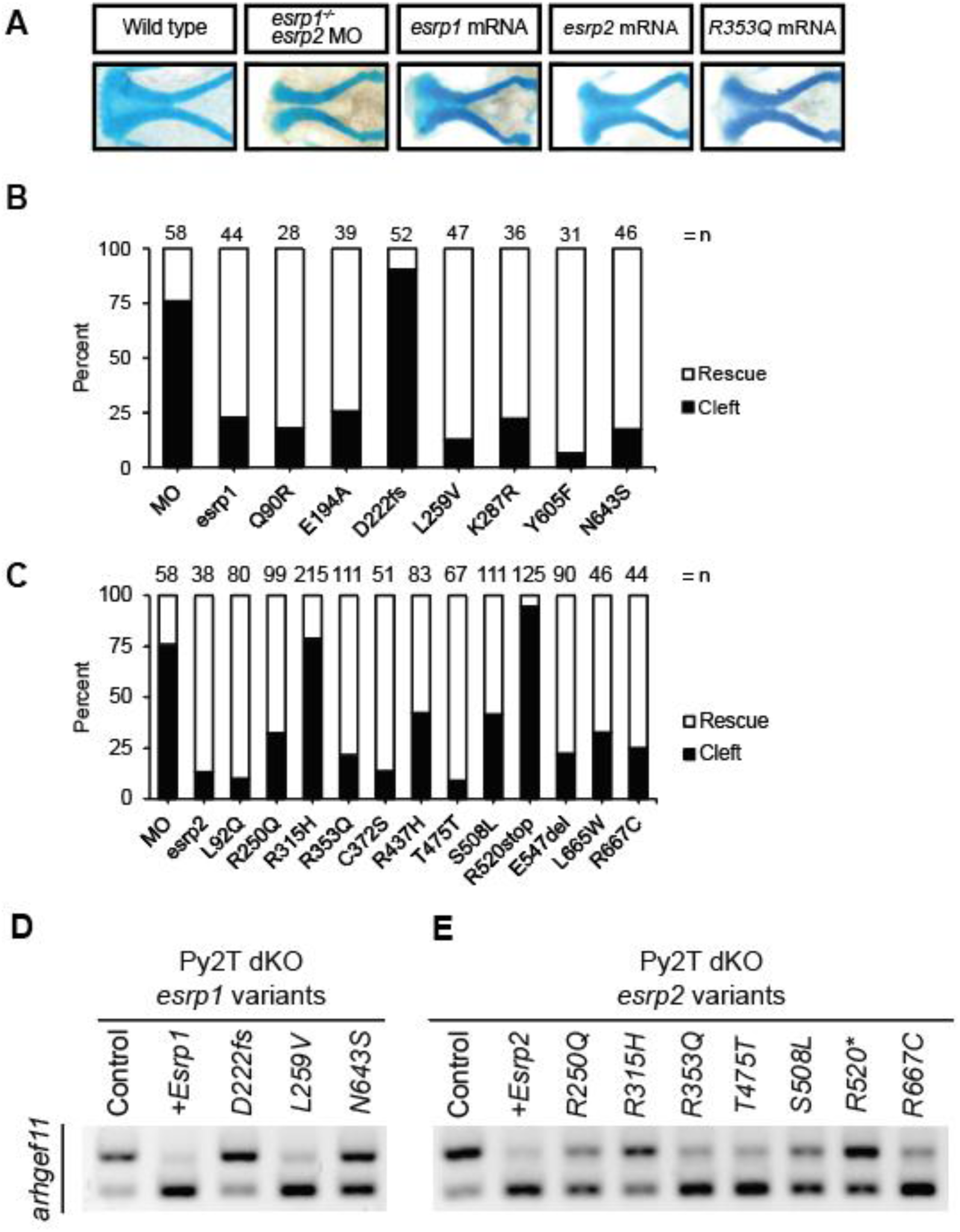
Functional testing of human ESRP1 and ESRP2 gene variants. **(A)** Representative images of the ANC from Alcian-blue stained larvae at 4 dpf after injection with *esrp2* MO and 200pg of: *esrp1* mRNA, *esrp2* R353Q mRNA. ANC was scored as a rescued ANC or cleft ANC **(B)** *ESRP1* and **(C)** *ESRP2* gene variant rescue assay results for embryos injected with *esrp2* MO and 200pg of *esrp1* variant mRNA. Results presented as percentage of rescue vs. cleft as different numbers of embryos survived and were analyzed, indicated as n above each bar. **(D)** *ESRP1* and **(E)** *ESRP2* gene variant rescue assay by detecting alternative splicing of Arhgef11 in murine Py2T wildtype and *Esrp1−/−; Esrp2−/−* double knockout cells.

To test for the ability of human *ESRP1/2* gene variants to rescue the cleft ANC phenotype in zebrafish, each of the 19 *ESRP1* or *ESRP2* gene variants was co-injected with e*srp2*-MO into *esrp1^−/−^* zebrafish embryos. At 4 dpf, the injected fish were fixed, stained with Alcian Blue, and analyzed. We found that for *ESRP1,* all 6 missense variants rescued the ANC phenotype. Only one variant, a frameshift mutation at the 222 aspartate residue (D222fs), had a large proportion of cleft ANC in the injected clutch compared to embryos injected with wildtype *esrp1* mRNA, and was scored as a pathogenic variant (Figure 3B). For *ESRP2*, 10 out of 12 tested gene variants rescued the ANC phenotype, in a ratio like the *esrp2* mRNA control and were scored as benign variants. The silent mutation T475T, served as an internal negative control and also scored as benign. The remaining 2 *ESRP2* gene variants (R315H and R520*) failed to rescue the ANC phenotype and were scored as pathogenic (Figure 3C).

To independently assess the gene variant functional testing results obtained from the zebrafish model, we tested 3 *ESRP1* and 8 *ESRP2* human gene variants using the mouse Py2T cell assay, with epithelial-specific RNA splicing of *Arhget11* as the readout (Figure 3D, 3E). We aimed to obtain an additional functional assessment for those gene variants testing results that contradicted *in silico* prediction. We performed site-directed mutagenesis to introduce the 11 gene variants, that were electroporated into *Esrp1/2* DKO PY2T cells and performed the RT-PCR assay 24 hours post-electroporation. We found that for *ESRP1,* gene variant L259V restored *Arhgef11* restriction to the epithelial isoform was scored as damaging for Polyphen-2 and Alpha missense and benign for SIFT (Figure 3, Table 1). The frameshift variant, D222fs, that was pathogenic in the in vivo assay was also pathogenic in this assay as it was unable to restore the epithelial isoform (Figure 3D, Table 1). Interestingly, the *ESRP1* gene variant N643S partially restored some of the splicing function of *Esrp1*, where both epithelial and mesenchymal *Arhgef11* isoforms were detected in a 1:1 ratio (Figure 3D). However, the same variant, N643S, in zebrafish rescued the phenotype. Statistical analysis for the Py2T rescue assay, can be found at Supplementary Figure 2. These results suggest that *ESRP1* N643S variant may be hypomorphic, or that *Arhgef11* is just one readout of *Esrp1* mRNA splicing activity. Because *Esrp1* shows position-dependent repression of exon splicing of *Arhgef11*, it is possible that some domains or regions may be required, or not, for some specific functions. It is possible that some splicing events may be differentially affected by mutations and there are other suggested functions of *Esrp1* in mRNA stabilization or post-transcriptional regulation that are accounted for in the zebrafish rescue assay (60).

For *ESRP2,* variants R250Q, R353Q and R667C rescued the molecular splicing of *Arhgef11* in the Py2T assay, (Figure 3E, Table 1). However, *ESRP2* gene variants R315H, S508L, and R520* failed to rescue deficient *Arhgef11* splicing in the Py2T assay and were scored as pathogenic, corroborating the pathogenic scoring from the zebrafish ANC rescue assay (Figure 3E, Table 1).

Overall, we found that the *in vivo* zebrafish ANC rescue assay and the *in vitro* Py2T splicing assays were largely concordant to determine pathogenicity of the *ESRP1* and *ESRP2* gene variants tested. PolyPhen-2 correctly predicted the effect of 8/18 (44.4%) tested gene variants, while SIFT correctly predicted the effect of 7/18 (38.8%) gene variants. When the predictions of both algorithms were concordant, they correctly predicted the consequence of 5 out of 7 (71.4%) gene variants on protein function (Table 1). The performance of concordant predictions was better for annotating benign variants where the algorithms correctly identified all four concordant benign variants with benign effects in both of our assays. Strikingly, the computational agreement incorrectly annotated 2 of 4 (50%) gene variants as pathogenic that had benign effects in both rescue assays. Ultimately, the algorithmic predictions were unable to determine half of the identified gene variants and greatly overestimated the prevalence of pathogenic variants (Table 1).

### AlphaMissense over-interpreted pathogenic variants

Recently a new gene variant analysis tool AlphaMissense was released and purported to improve variant calling accuracy by leveraging protein structure information predicted by machine learning algorithm AlphaFold (72). Using AlphaMissense to analyze the 6 *ESRP1* and 9 *ESRP2* missense variants we had functionally tested, we observed that AlphaMissense classified 5 variants as benign for *ESRP1* (Q90R, E194A, K287R, Y605F, N643S) consistent with the functional tests, but called L259V as pathogenic when both the *in vivo* and *in vitro* functional tests demonstrated protein function (Figure 4).

**Figure 4.**
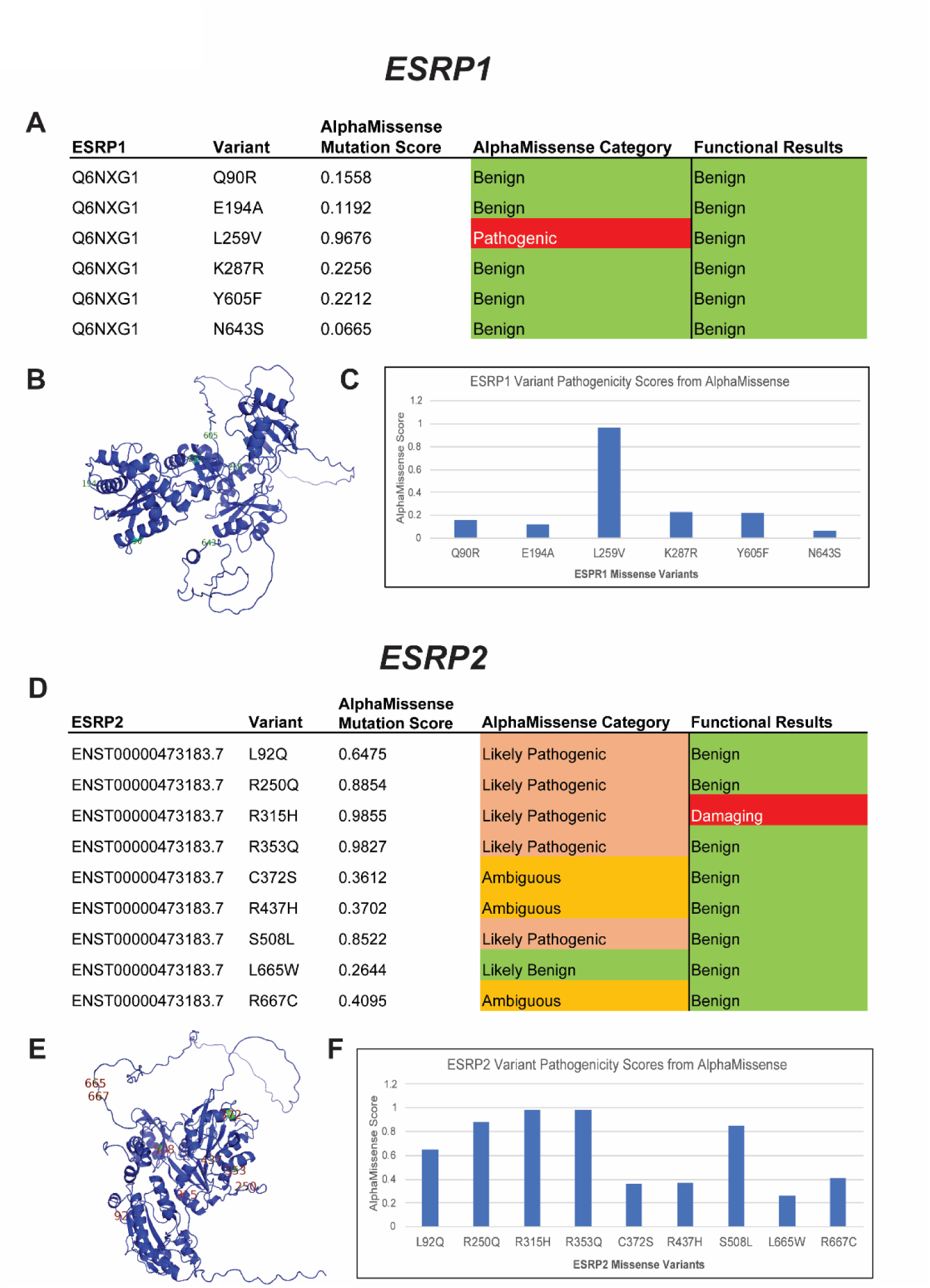
AlphaMissense pathogenicity predictions for ESRP1 and ESRP2 missense variants. *ESRP1* and *ESRP2* gene variants from OFC cases in the GMFK Children’s dataset and ClinVar variants associated with cleft lip and/or palate or autosomal recessive deafness were identified. 6 ESRP1 and 9 ESRP2 (A and D) missense variants were analyzed using the AlphaMissense (AM) model. The tables (C and F) show the AM-predicted pathogenicity compared to our functional test results and the AM mutation score, which is also graphed. On the left, the *ESRP1* and *ESRP2* AlphaFold structures (B and E), with labeled missense mutations, color-coded with the functional results.

For *ESRP2*, AlphaMissense and the experimental validation were only concordant on 2 variants out of 9, calling R315H as pathogenic and L665W as benign (Figure 4). AlphaMissense called 6 variants as pathogenic when they were shown to be functionally benign in both *in vitro* and *in vivo* functional tests. Therefore, our results showed that AlphaMissense may over-interpret variants as pathogenic for some genes.

### Alternative splicing to generate epithelial isoform of Ctnnd1 requires Esrp1/2 function

We and others demonstrated that Esrp1 and Esrp2 regulate the alternative splicing of Ctnnd1, generating isoforms that differ between epithelial and mesenchymal cell types (20, 60, 66, 73), making *Ctnnd1* an interesting *Esrp1/2* target that has also been implicated in CL/P (61).

*CTNND1* (p120-catenin) have been associated with Blepharocheilodontic (BCD) syndrome and non-syndromic human CL/P (19, 20, 74). Like other catenins, *Ctnnd1* has dual roles: it functions as part of the adherens junction cellular scaffolding to stabilize cell adhesion molecules, as well as a transcriptional regulator (19, 75–79). Furthermore, functional differences between epithelial and mesenchymal forms of *Ctnnd1* have been described (80–82). Four major isoforms for *Ctnnd1* have been characterized in humans. The full-length isoform, isoform 1, has a translational start site at the first methionine in the sequence (1 Met), while isoforms 2, 3, and 4 undergo splicing events that cause a 5’ truncation of the transcript and change the translational start site to methionines 55, 102, and 324, respectively. Isoform 1 of *CTNND1* is predominantly expressed in the mesenchyme, while the shorter isoform 3 is restricted to the epithelium. The remaining isoforms, 2 and 4, are less abundant and have not been thoroughly characterized (74).

When we aligned the amino acid sequences between human, mouse, and zebrafish *Ctnnd1* homologs, we found that methionine in positions 1 and 102 are conserved in all three species. Methionine 55 is part of a 14 aa stretch absent in zebrafish (Figure 5A). Given that transcripts for the long (mesenchymal) isoform shifts to the shorter (epithelial) isoform by splicing out a 5’ exon(s) and moving down to a conserved methionine, splicing pattens are well-conserved across human, mouse, and zebrafish. Cox et. al reported that *ESRP2* and a short form of the full-length *CTNND1* protein, identified by an antibody to the C-terminus, are colocalized in the periderm of human embryos (20). Meanwhile, RNA splicing of *Ctnnd1* transcripts is deficient in the embryonic epithelium of *Esrp1^−/−^* mice (53).

**Figure 5.**
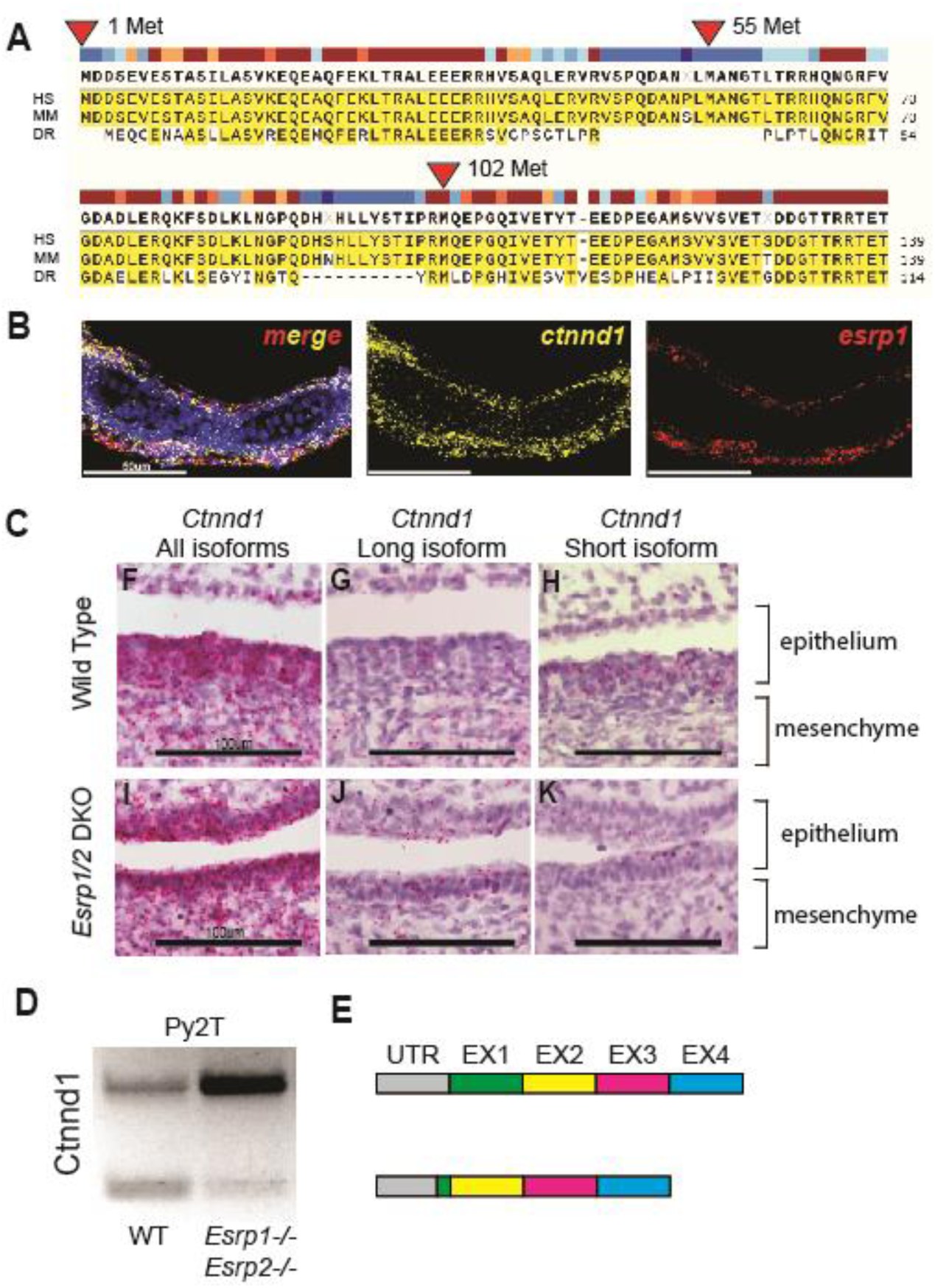
Alternative splicing of *Ctnnd1* is regulated by Esrp1/2. **(A)** Amino acid sequence alignment of the first 140 residues of CTNND1 protein across human, mouse, and zebrafish. Translation for isoform 1 of CTNND1 begins at methionine 1, while isoform 3 encodes a truncated form that starts translation at methionine 102. Methionine residues at positions 55 and 324 are not conserved across all three species. **(B)** Detection of *esrp1* and *ctnnd1* gene expression in zebrafish at 4dpf, demonstrates shared localization of transcripts in the embryonic epithelium. This coronal section includes the ventral Meckel’s cartilage. **(C)** Detection of murine *Ctnnd1* mRNA using isoform-specific base-scope probes in the oral epithelium and tongue mesenchyme. The wildtype sections show that the *Ctnnd1* long isoform is present in both epithelial and mesenchymal cells. The *Ctnnd1* short isoform is present preferentially in epithelial cells and not in the mesenchymal cells. In the *Esrp1/2* DKO mouse, the mesenchymal *Ctnnd1* long isoform is detected in epithelial and mesenchymal cells, with loss of the *Ctnnd1* short isoform. **(D)** RT-PCR of the *Ctnnd1* long and short isoforms from Py2T cells. **(E)** Diagrammatic representation of the ESRP-regulated *CTNND1* alternative splicing to generate the shorter epithelial isoform.

We confirmed that long and shorter *Ctnnd1* isoforms were found in the mouse Py2T cells by performing RT-PCR using primers spanning exon 2, which is partially skipped in the shorter isoform for *Ctnnd1*. In the *Esrp1/2*−/− Py2T cell line, the splicing pattern of *Ctnnd1* shifts and is biased towards the longer mesenchymal isoform, confirming previous observations (60).

To localize *Ctnnd1* and *Esrp1/2* gene expression in wildtype mouse and zebrafish, we carried out RNAscope and BaseScope on wildtype and mutant mouse and zebrafish sections (Figure 5B 5C). The *Ctnnd1* probe used identifies shared C-terminal exons shared in all *Ctnnd1* isoforms. Only *Esrp1* probe was used here as we and others have previously shown that *Esrp1* and *Esrp2* gene expression are co-localized in mouse and zebrafish (21, 22, 52, 66). In zebrafish, *ctnnd1* and *esrp1* RNAscope signals are co-localized robustly throughout the oral epithelium with sparse signals in the mesenchyme.

To assess the tissue specific distribution of the longer mesenchymal isoforms of *Ctnnd1* vs. shorter epithelial isoform, BaseScope probes were used to detect the two *Ctnnd1* isoforms from wildtype and *Esrp1*−/−; *Esrp2−/−* mutant mouse at E15. Similar to RNAScope result in zebrafish (Figure 5B), the murine *Ctnnd1* BaseScope signals for both mesenchymal and epithelial isoforms were robust in the oral epithelium and sparsely scattered in the mesenchyme (Figure 5D and 5G). When signal is differentiated by isoform, the longer *Ctnnd1* mesenchymal isoform was uniformly distributed throughout the epithelium and mesenchyme (5E and 5H). However, the shorter *Ctnnd1* epithelial isoform was restricted to the epithelial cells and excluded from the muscle (5F and 5I). In the wildtype, BaseScope signals of the longer *Ctnnd1* mesenchymal isoform appeared equally distributed in the mesenchyme and epithelium, and the signals of the shorter isoform was epithelial restricted. In the *Esrp1−/−; Esrp2−/−* mutant mouse, *Ctnnd1* transcript level was significantly reduced and predominantly the longer *Ctnnd1* mesenchymal isoform was detected, in both the mesenchyme and epithelium. The shorter *Ctnnd1* epithelial isoform was sparsely detected via BaseScope in the *Esrp1−/−; Esrp2−/−* mutant, consistent with the finding where shorter isoform was significantly reduced in the *Esrp1−/−; Esrp2−/−* Py2T cells by qPCR. These results corroborate that Esrp1/2 is required for RNA splicing of *Ctnnd1*, generating the shorter isoform specifically in the epithelium but not the mesenchyme.

### *CTNND1* gene variants from OFC cohorts

Twenty-four *CTNND1* gene variants have been reported and a growing number of new variants have been found in ongoing WGS studies of OFC cohorts (19, 26). In a recent analysis of 759 OFC trios, we identified 15 variants in *CTNND1* with allele frequencies less than 0.1% in gnomAD (Figure 3 Supplementary Material). Two variants were *de novo* and one was inherited from an affected parent. Pathogenic variants in *CTNND1* accounted for 0.8% of the cohort. Only 10% of the cohort had a pathogenic variant in 500 genes implicated in OFC that we analyzed, making *CTNND1* the mostly frequently mutated variant in this cohort (61). In the gene-based burden test, rare variants were nominally over-transmitted to affected children (p=0.06); *de novo* variants are enriched in *CTNND1* (p=0.005 for loss-of-function *de novo* variants; 0.001 for protein-altering *de novo* variants). Nearly all the missense variants were classified as variants of unknown significance, indicating that functional testing is critical. In fact, we estimate that *CTNND1* mutations account for at least 1.5% of CL/P cases. By comparison, *IRF6* mutations are estimated to be the most common cause of CL/P, accounting for 2% of cases. Taken together, *CTNND1* stands to be as important as *IRF6* in contributing to the genetic risk of syndromic and non-syndromic CL/P.

### *Ctnnd1* over-expression rescue *esrp 1−/−; esrp 2−/−* cleft ANC, curled fin and fused otolith phenotypes

To functionally assess the relationship between *Esrp* and *Ctnnd1*, we injected the zebrafish *ctnnd1* isoform-201 (ENSDART00000106048.4) mRNA into *esrp1*^−/−^; *esrp2*^+/−^ offspring at the 1-cell stage. Mutants and control embryos were analyzed at 4 dpf, assessing the ANC, the pectoral fin, and otoliths phenotypes, followed by genotyping (figure 6A).

**Figure 6.**
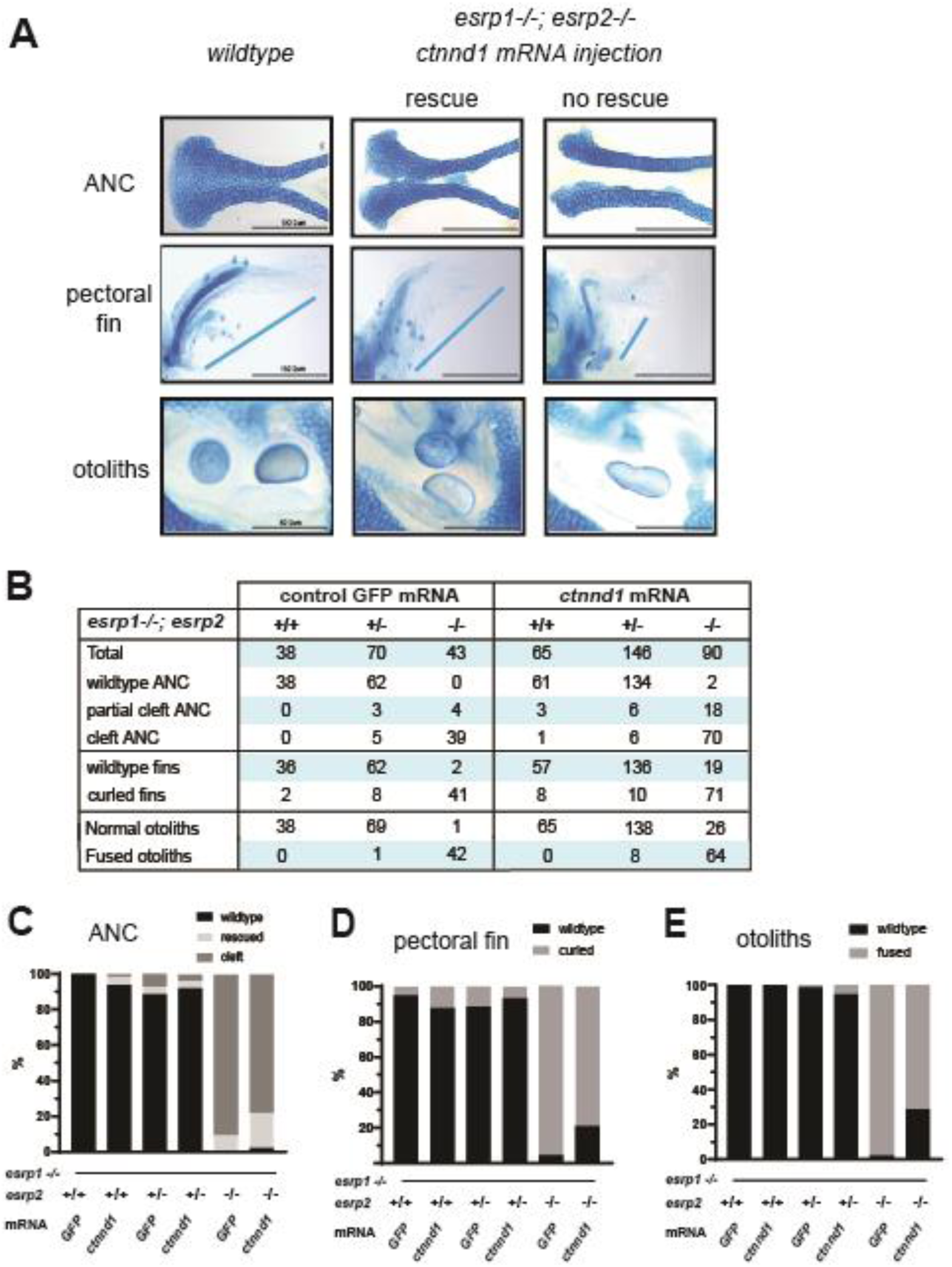
Over-expression of *ctnnd1* rescues esrp1^(−/−)^, esrp2^(−/−)^ epithelial phenotypes. **(A)** Image representing how wild type, intermediate and cleft ANC, pectoral fins and otoliths were sorted. **(B)** Representative table with the number of total fish injected and rescued by the *ctnnd1* mRNA injection with GFP mRNA injection as control. Scoring of ANC phenotype (%) **(C)**, fin phenotype (%) **(D)** and the otolith phenotype (%) **(E)** in the injected *esrp1−/−; esrp2+/−* inter-cross larvae confirmed by genotyping, showing 20-22% rescue of ANC, fin and otolith phenotypes in the *esrp1−/−; esrp2−/−* double homozygous larvae.

Control *gfp* mRNA injected *esrp1−/−; esrp2−/−* larvae, exhibited cleft ANC, the pectoral fins were hypoplastic and stuck to the thorax, and fused otoliths, the mutant phenotypes were fully penetrant and reliably scored (Figure 6B-E). In the *ctnnd1* mRNA injected *esrp1−/−; esrp2−/−* larvae, 22% (n = 20 of 90, p<0.01) demonstrated a full or partial rescue of the ANC (Figure 6B-E). Correspondingly, the injected *esrp1−/−; esrp2−/−* larvae exhibited significant rescue of the abrogated fin phenotype, with 21% (n = 19 of 90, p<0.01) exhibiting extension of the pectoral fin and angling away from the thorax. The fused otolith phenotype was scored as either separate or fused, and demonstrated 26% (n = 26 of 90, p<0.01) rescue (Figure 6A). The morphogenesis of the ANC, pectoral fin and the otoliths all reflect different aspects of embryonic epithelium development and interaction with the associated mesenchyme of the *esrp 1−/−; esrp 2−/−* embryos. The *ctnnd1* mRNA over-expression rescuing the epithelial defects in the *esrp 1−/−; esrp 2−/−* suggests that a key function of *esrp1/2* in epithelial biology is to regulate *ctnnd1* function.

## Discussion

Several independent lines of evidence corroborate that the *ESRP1* and *ESRP2* genes are important OFC loci in humans. *ESRP1* was proposed to be the most likely candidate CL/P risk gene in the 8q22.1 locus (83, 84). Ectopic expression of p63 converted human fibroblasts to keratinocyte-like cells and *ESRP1* was transcriptionally induced together with activation of an epithelial enhancer within a topologically associated domains (TADs) containing a non-syndromic CL/P risk locus (85). This is consistent with the biological observation and *p63*, *Irf6* and *Esrp1/2* co-localize in the embryonic epithelium, and that mutations of these 3 genes result in OFC phenotypes. Further, a whole exome sequencing study of non-syndromic CL/P in multi-affected families identified pathogenic variants in *ESRP2* with an autosomal dominant inheritance pattern (20).

Several studies showed in mouse and zebrafish models that *Esrp1* and *Esrp2* are important in craniofacial development. We showed that *Esrp1* and *Esrp2* are co-localized with *Irf6* in the embryonic oral epithelium, and when *Ersp1/2* are disrupted, cleft of the lip and palate formed, validating that mouse and zebrafish are robust animal models of human OFC (21, 52, 53).

There is growing recognition that RNA binding proteins that regulate alternative splicing play vital roles in craniofacial morphogenesis. Clinically, spliceosomopathies are often associated with syndromic craniofacial abnormalities due to disruption of splicing factors such as *PUF60*, *ETUD2*, *SF3B4*, *RBM10*, and *ESRP2* (86). Animal models defective in RNA splicing that exhibit craniofacial phenotypes include: *Esrp1/2, Rbfox2*, *Srsf3*, and *Sf3b2* (21, 22, 52, 87, 88). The ESRP proteins are uniquely expressed in epithelial structures and direct post-transcriptional modifications that distinguish protein isoforms between epithelium and mesenchyme. We applied complementary phenotypic and molecular assays to interrogate the functional consequence of identified *ESRP1/2* gene variants in cohorts of autosomal recessive deafness and CL/P.

As the magnitude of available WGS data increases, the need for assigning clinically actionable information continues to grow. The sequence variant interpretation (SVI) working group from ACMG-AMP frequently reconvenes to update, revise, and refine the ACMG criteria to provide the clearest guidance possible (33, 34). Most recently, the working group provided further guidance regarding functional assays and experimental model systems. Among these, they highlighted the need to ascertain the gene variants’ physiologic context and molecular consequence. Here, we applied complementary phenotypic assays in the zebrafish ANC rescue, in addition to the Py2T splicing assay, to assess the physiologic and molecular consequences of *ESRP1/2* gene variants observed in clinical cohorts. These functional tests identified 7 pathogenic variants out of 18 *ESRP1/*2 variants examined. Moreover, these functional readouts of orthologous systems across species attest to the strongly conserved nature of epithelial splicing by the *ESRPs* in craniofacial morphogenesis. These results highlight the need for experimental models to enhance the validity of *in silico* predictions of protein function. We found that while the SIFT and PolyPhen-2 algorithms have a positive predictive value when they align in predicting benign variants, they tend to overestimate the prevalence of pathogenic variants.

While AlphaMissense provided slightly better predictions for *ESRP1* than SIFT and PolyPhen-2, in the case of *ESRP2*, AlphaMissense over-interpreted benign variants as pathogenic. A similar high false positive rate was seen in a different disease, cystic fibrosis transmembrane conductance regulator (89), and for epithelial master regulator *IRF6* (90). This work highlights that protein structure and machine learning approaches today are still insufficient to accurately predict pathogenicity, where functional tests are indispensable to validate the pathogenicity of variants.

These functional assays revealed novel insights into *ESRP1/2* protein function and downstream targets spliced by the *ESRP*s. We found that the gene variants with the largest effect size for the zebrafish ANC rescue assay lie in RRM1 and RRM3 of *ESRP2.* Variants R250Q and R353Q were predicted by PolyPhen-2, SIFT and AlphaMissense to be damaging or likely pathogenic, but in both independent functional tests corroborated to be benign variants. In contrast, R315H was functionally tested by both assays to be a deleterious variant, consistent with prior work demonstrating R315 to impact RNA binding based on protein structure analysis (91). Furthermore, we provide molecular evidence that *Esrp* transcripts rescue molecular splicing patterns of putative *Esrp-*target genes *Arhgef11* and *Ctnnd1*. Moreover, gene variants with pathogenic potential do not restore splicing patterns of *Arhgef11*, providing evidence that the gene variants impair *Esrp* function and likely contribute to disease pathogenicity. These functional assays provide key data to satisfy the ACMG-AMP standards, where molecular assays are used to contribute to our understanding of mechanisms for disease.

Mutations in *CTNND1* and *CDH1* (E-cadherin) are the known cause of BCD, which includes abnormal eyelids, upper lip, palate, and teeth development (20, 74, 92). The precise pathological mechanism remains to be elucidated, but in healthy epithelial cells *CTNND1* binds to E-cadherin to stabilize adherens junctions and desmosomes, and therefore displacement of *CTNND1* causes endocytosis of *CDH1* and loss of the junction. Another possibility is disruption of the canonical WNT pathway signaling, as *CTNND1* is known to modulate transcription by binding to transcription factors such as Kaiso in the *Wnt* pathway (93, 94). It is known, and further supported by the evidence in this work, that alternatively spliced isoforms of *CTNND1* are differentially expressed in the epithelium and mesenchyme, and here we show that those distinct splicing patterns are dependent on *Esrp1/2* activity. However, it is not known how the alternatively spliced isoforms differ in function, alter embryonic and craniofacial morphogenesis, or contribute to disease. Thus, further studies into the functional differences between *CTNND1* isoforms are warranted and would provide insight into the disease etiology of BCD or the mechanism of the cleft palate from *ESRP* loss-of-function.

## Acknowledgements

We thank the Aquatics Core at Massachusetts General Hospital and Children’s Hospital of Philadelphia (CHOP) for their dedication to fish health and maintenance of our colonies. We thank the CHOP IDDRC Biostatistics and Data Science core (HD105354) for consultation. We are grateful for the funding support from the National Institutes of Health (HG013031) to K.W. and (DE032332, DE027983) to E.C.L., research and fellowship grants from Shriners Hospitals for Children and institutional support from Children’s Hospital of Philadelphia.

## Notes

### Competing Interest Statement

The authors have declared no competing interest.

