## Supplemental information for "Functional analysis of *ESRP1/2* gene variants and *CTNND1* isoforms in orofacial cleft pathogenesis"

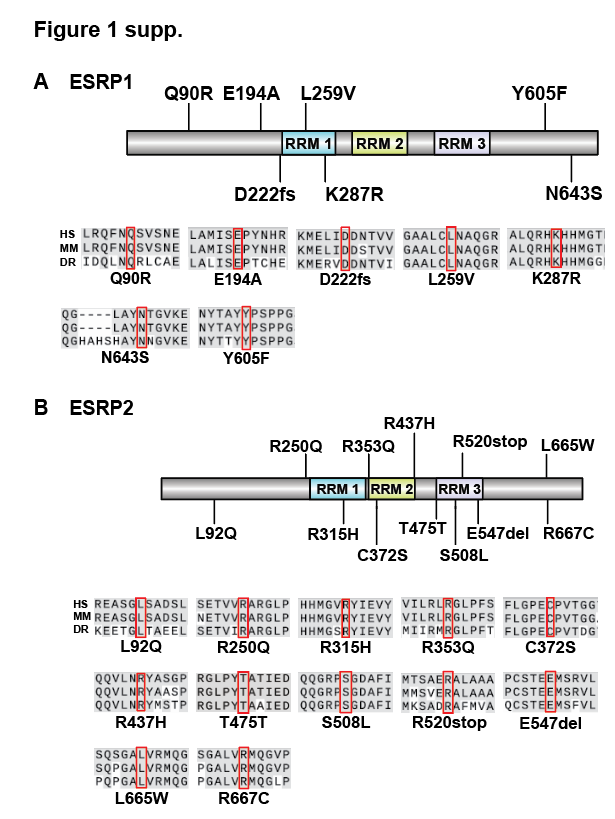
Supplemental Information

**Supplementary Figure 1. Selection of evolutionarily conserved *ESRP1* and *ESRP2* gene variants.** *ESRP1* and *ESRP2* gene variants were identified from OFC cases in the GMFK Children’s dataset and ClinVar variants associated with cleft lip and/or palate or autosomal recessive deafness. Gene variants disrupting amino acid residues that are fully conserved between humans, mice, and zebrafish were selected: 7 in *ESRP1*, 12 in *ESRP2*. **(A)** Schematic of the ESRP1 protein labeled with 7 identified gene variants, including two variants in RRM1. Truncated alignments surrounding the gene variants are provided below the diagram. **(B)** Schematic of the ESRP2 protein labeled with 12 identified gene variants, including two variants in each of RRM1 and RRM2, three variants are in RRM3. A silent variant *ESRP2* T475T was used as a negative control. Truncated alignments surrounding the gene variants are provided below the diagram. Fully conserved residues between human, zebrafish, and mouse amino acid sequences surrounding the gene variants are highlighted in grey.

**Figure 2 supp.**

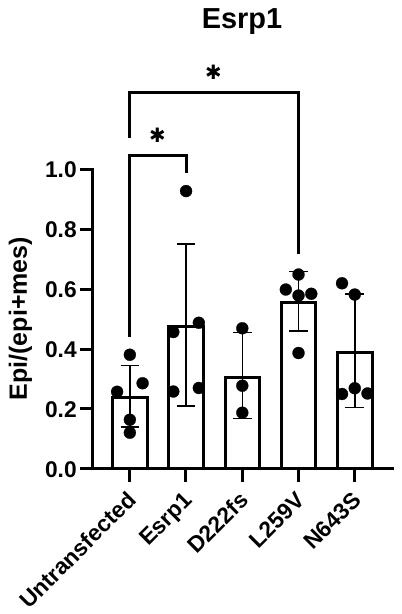

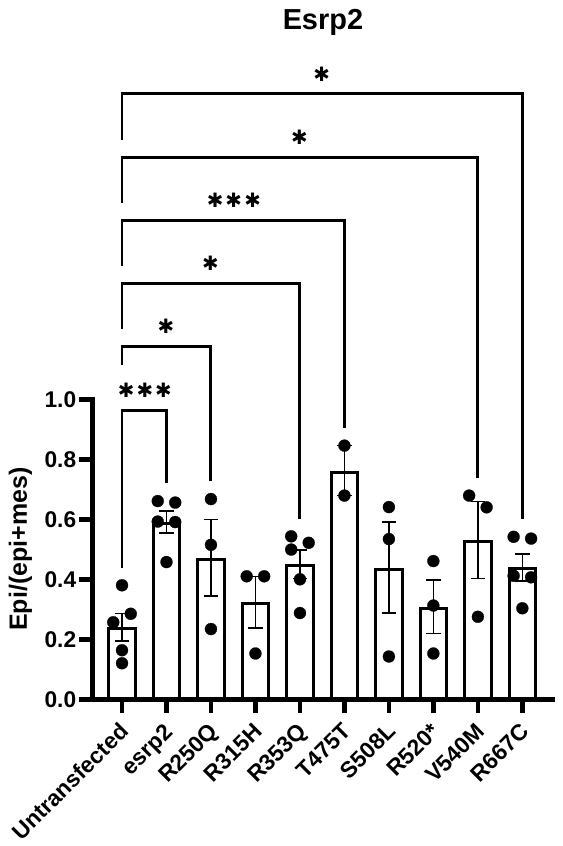

**Supplementary Figure 2. Py2T rescue assay statistical analysis.** *ESRP1* and *ESRP2* specific variants were analyzed using ordinary one-way ANOVA to confirm rescue statistical significance in comparison with untransfected cells. Data mean ± SEM, *p<0.05, ***p<0.001.

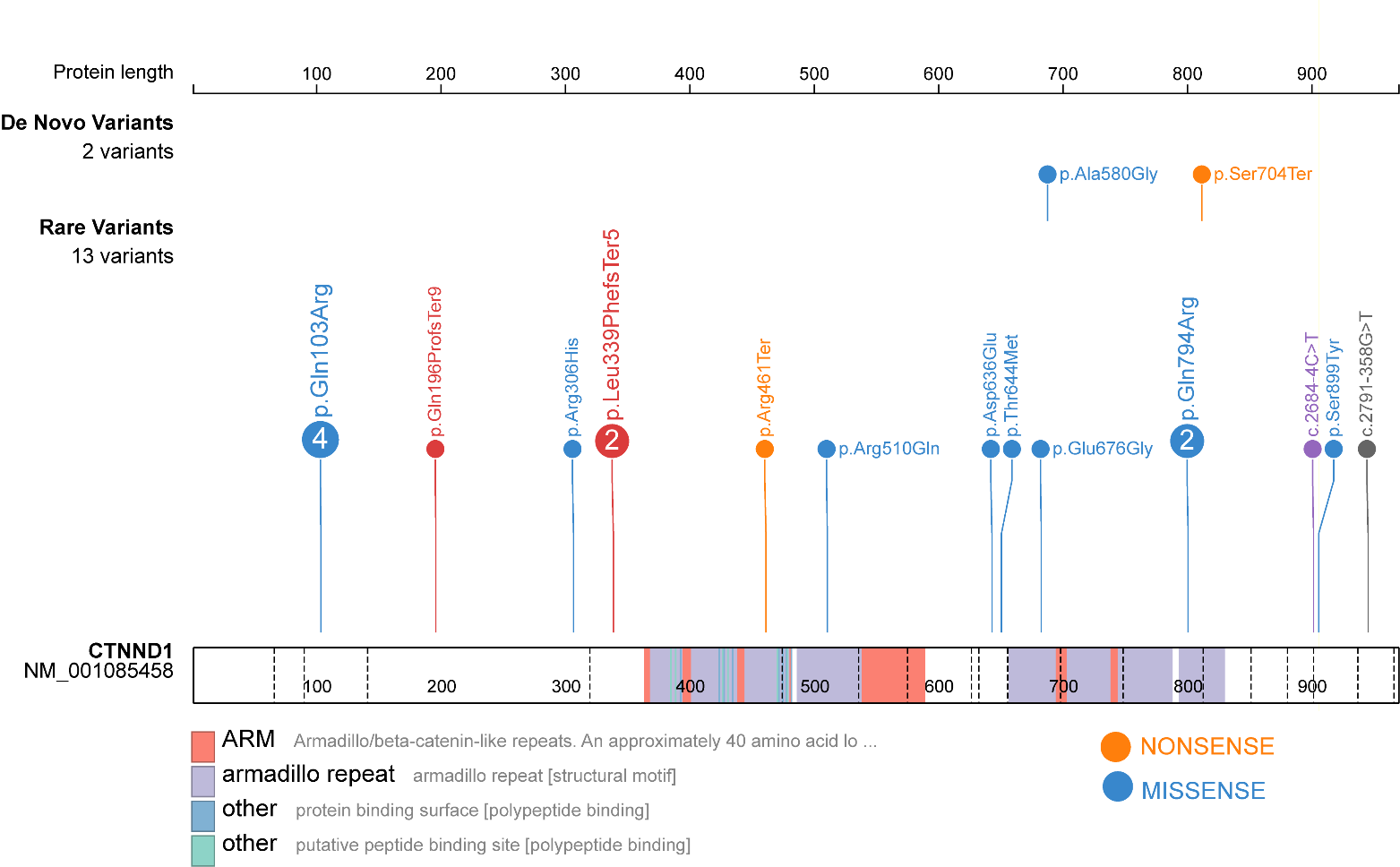
**Figure 3 supp.**

**Supplementary Figure 3. CTNND1 variants from OFC cohorts.** 15 variants were identified in 759 OFC trios. 2 Variants were de novo, and 13 were considered rare variants.

Supplemental Table 1

| ESRP1 | | | | | | | | |
| --- | --- | --- | --- | --- | --- | --- | --- | --- |
| variant | **CHR:BP (hg38)** | **Ref Allele** | **Alt Allele** | **rs id** | **Transcript Change** | **AA Change** | **Source** | **Associated disease** |
| Q90R | 8:94643310 | A | G | rs778878601 | NM_017697.4:c.269A>G | NP_060167.2:p.Gln90Arg | GMKF | - |
| E194A | 8:94662362 | A | C | rs1302067900 | NM_017697.4:c.581A>C | NP_060167.2:p.Glu194Ala | GMKF | - |
| D222fs | 8:94664716-94664734  -19bp deletion bp 665-683 | - | - | - | NM_017697.4:c.665_683del | NP_060167.2:p.Asp222fs | ClinVar | Hearing loss,  autosomal recessive 109 |
| L259V | 8:94664946 | C | G | rs1554577402 | NM_017697.4:c.775C>G | NP_060167.2:p.Leu259Val | ClinVar | Hearing loss,  autosomal recessive 109 |
| K287R | 8:94665031 | A | G | rs747729420 | NM_017697.4:c.860A>G | NP_060167.2:p.Lys287Arg | GMKF | - |
| Y605F | 8:94678365 | A | T | rs1228296571 | NM_017697.4:c.1814A>T | NP_060167.2:p.Tyr605Phe | GMKF | - |
| N643S | 8:94692784 | A | G | rs767362823 | NM_017697.4:c.1928A>G | NP_060167.2:p.Asn643Ser | GMKF | - |

| ESRP2 | | | | | | | | |
| --- | --- | --- | --- | --- | --- | --- | --- | --- |
| variant | **CHR:BP (hg38)** | **Ref Allele** | **Alt Allele** | **rs id** | **Transcript Change** | **AA Change** | **Source** | **Associated disease** |
| L92Q | 16:68235686 | A | T | rs1567566981 | NM_024939.3:c.275T>A | NP_079215.2:p.Leu92Gln | GMKF | - |
| R250Q | 16:68232649 | C | T | rs755729355 | NM_024939.3:c.749G>A | NP_079215.2:p.Arg250Gln | ClinVar | Cleft lip with or without cleft palate |
| R315H | 16:68232381 | G | A | rs751873605 | NM_024939.3:c.944G>A | NP_079215.2:p.Arg315His | ClinVar | Cleft lip with or without cleft palate |
| R353Q | 16:68232043 | C | T | rs201908706 | NM_024939.3:c.1058G>A | NP_079215.2:p.Arg353Gln | GMKF | - |
| C372S | 16:68231987 | A | T | rs149234558 | NM_024939.3:c.1114T>A | NP_079215.2:p.Cys372Ser | GMKF | - |
| R437H | 16:68231684 | C | T | rs750375203 | NM_024939.3:c.1310G>A | NP_079215.2:p.Arg437His | GMKF | - |
| T475T | 16:68231569 | G | A | rs56323755 | NM_024939.3:c.1425G>A | NP_079215.2:p.Thr475= | GMKF | - |
| S508L | 16:68231366 | C | T | rs143677348 | NM_024939.3:c.1523C>T | NP_079215.2:p.Ser508Leu | ClinVar | Cleft lip with or without cleft palate |
| R520stop | 16:68231331 | C | T | rs142168544 | NM_024939.3:c.1558C>T | NP_079215.2:p.Arg520Ter | ClinVar | Cleft lip with or without cleft palate |
| E547del | 16: 68231248-68231250 | GAG | - | - | NM_024939.3:c.1636GAG | NP_079215.2:p.Glu547del | ClinVar | Cleft lip with or without cleft palate |
| L665W | 16:68230459 | A | C | rs373594090 | NM_024939.3:c.1994T>G | NP_079215.2:p.Leu665Trp | GMKF | - |
| R667C | 16:68230454 | G | A | rs575114143 | NM_024939.3:c.1999C>T | NP_079215.2:p.Arg667Cys | GMKF | - |
